# Tumor cell-intrinsic HFE drives glioblastoma growth

**DOI:** 10.1101/2022.04.13.487917

**Authors:** Katie M. Troike, Daniel J. Silver, Prabar K. Ghosh, Erin E. Mulkearns-Hubert, Christopher G. Hubert, James R. Connor, Paul L. Fox, Bjarne W. Kristensen, Justin D. Lathia

## Abstract

**Background:** Glioblastoma (GBM) tumor cells modulate expression of iron-associated genes to enhance iron uptake from the surrounding microenvironment, driving proliferation and tumor growth. The homeostatic iron regulator (*HFE*) gene, encoding the iron sensing HFE protein, is upregulated in GBM and correlates with poor survival outcomes. However, the molecular mechanisms underlying these observations remain unclear. Identification of pathways for targeting iron dependence in GBM tumors is therefore a critical area of investigation.

**Methods:** We interrogated the impact of cell-intrinsic *Hfe* expression on proliferation and tumor growth through genetic loss and gain of function approaches in syngeneic mouse glioma models. We determined the expression of iron-associated genes and their relationship with survival in GBM using public datasets and identified differentially expressed pathways in *Hfe* knockdown cells through Nanostring transcriptional profiling.

**Results:** Loss of *Hfe* induced apoptotic cell death *in vitro* and inhibited tumor growth *in vivo* while overexpression of *Hfe* accelerated both proliferation and tumor growth. Analysis of iron gene signatures in *Hfe* knockdown cells revealed alterations in the expression of several iron-associated genes, suggesting global disruption of intracellular iron homeostasis. Analyzing differentially expressed pathways further identified oxidative stress as the top pathway upregulated with *Hfe* loss. Enhanced ^55^Fe uptake and generation of reactive oxygen species (ROS) were found with *Hfe* knockdown, implicating toxic iron overload resulting in apoptotic cell death.

**Conclusions:** Collectively, these findings identify a novel role for HFE in regulating iron homeostasis in GBM tumors and provide a potential avenue for future therapeutic development.

**Key Points:**

- HFE is an iron sensor that is upregulated in GBM and negatively impacts survival.
- HFE overexpression drives proliferation and tumor growth *in vivo*.
- Loss of HFE increases production of reactive oxygen species and induces apoptosis, extending survival *in vivo*.

**Importance of Study:** Dysregulation of iron metabolism is an important feature of GBM contributing to tumor growth and negatively impacting survival. The identification of key iron regulators controlling this process is therefore important for therapeutic targeting. We identify HFE as an important regulator of iron homeostasis in GBM and suggest a role for sexual dimorphism in HFE-mediated tumor iron regulation that ultimately results in differential survival outcomes. Our findings demonstrate that HFE drives tumor cell proliferation and survival in GBM and may be a viable target for modulating tumor iron flux and inducing apoptosis in tumor cells.

## Introduction

Glioblastoma (GBM) is the most common primary malignant brain tumor and has an exceptionally poor prognosis. When treated with standard-of-care therapy, which includes maximal safe surgical resection, radiation, and chemotherapy, the median survival for patients with GBM is approximately 20 months^1^. Multiple factors have been identified that facilitate GBM growth and therapeutic resistance, culminating in disease recurrence. Tumor cell-intrinsic factors, including cellular heterogeneity, metabolic, and genetic aberrations, along with extrinsic factors within the neural, endothelial, and immune cell compartments, drive tumor progression and contribute to therapeutic failure^2^. Recently, patient sex has been reported as a determinant of incidence and survival in GBM, with male patients exhibiting greater risk of developing GBM, as well as worse prognosis, when compared to female patients^3^. Sex differences are mediated through the effects of sex chromosomes and sex hormones, which influence a variety of processes in GBM, including malignant transformation, epigenetic landscape, anti-tumor immunity, and therapeutic response^4–6^. As such, the intersection of sex and other tumor cell-intrinsic and cell-extrinsic features is a critical area of investigation. One biological process known to exhibit a large degree of sexual dimorphism and to drive radioresistance and tumor progression in GBM is iron metabolism, an attractive focus for therapeutic development^7,8^.

Iron is an indispensable element required for enzymatic function in a number of normal cellular processes such as ATP production, DNA synthesis, and cell cycle regulation^9^. The brain is dependent on iron for normal development and function, playing a role in neurotransmitter synthesis, myelination, and microglia polarization^10^. The redox cycling potential of iron contributes to its pro-tumorigenic effects, including the generation of free radicals, which can damage DNA and promote malignant transformation^9^. Therefore, a complex regulatory mechanism involving iron uptake, storage, and release, mediated by a variety of iron-handling proteins, ensures the tight maintenance of intracellular iron levels to prevent toxicity. Dysregulation of iron metabolism, primarily driven by the aberrant expression of iron-handling proteins, is a hallmark of the tumor state, and increased import accompanied by reduced export is frequently observed in many different cancers^11^. Iron is therefore a putative therapeutic target for cancer, although the use of iron chelators *in vivo* has been limited by their lack of tumor specificity and their side effects^12^. GBM tumors exhibit increased iron uptake compared to non-tumor brain tissue, a property that has been exploited for specific targeting and imaging of these lesions^13,14^. Previous work has demonstrated that cancer stem-like cells within GBM tumors are efficient iron scavengers, upregulating their expression of specific iron-handling proteins to enhance uptake and storage and drive proliferation^15^. Conversely, disrupting intracellular iron storage in GBM cells induces cell death by several different mechanisms, including iron-dependent ferroptosis^16^. Given the pivotal role of iron in tumor initiation and growth, further insight into its regulation in GBM is necessary to improve understanding of this disease and to aid in the development of new therapies.

In a normal cellular state, iron homeostasis is maintained through a tightly regulated balance of iron import, export, and storage. When this balance is disrupted, iron can accumulate and lead to pathologies such as hereditary hemochromatosis (HH), an iron overload disorder. Dysregulated iron absorption in HH causes tissue iron accumulation, culminating in oxidative damage and cell death^17^. Additionally, ferroptosis, an iron-dependent form of cell death, has been reported to contribute to tissue damage in mouse models of hemochromatosis^18^. Most cases of HH are driven by mutations in the homeostatic iron regulator (*HFE*) gene, which encodes the transmembrane HFE protein^19^. In normal cells, HFE acts as a cellular iron sensor, mediating the uptake and release of iron indirectly through interactions with other iron-associated proteins^20^. Prior work in GBM has demonstrated an inverse relationship between HFE expression and survival in female patients^21^. However, no study to date has clarified how HFE expression modulates GBM patient survival at the molecular mechanistic level. Here, we demonstrate that *Hfe* loss in glioma cells enhances iron uptake and the generation of reactive oxygen species, which in turn promotes tumor cell death and extends subject survival. These findings underscore the importance of HFE in iron homeostatic maintenance and its potential as a target for therapeutic management of GBM.

## Results

### HFE is upregulated in GBM tumors and is associated with worse survival in female patients

Aberrant iron metabolism is considered a hallmark of the tumor state and can be leveraged by tumor cells to fuel their growth. We confirmed the presence of iron in patient GBM samples using Perl’s Prussian blue stain (**Figure 1A**). To validate the functional effect of iron depletion and supplementation on tumor cell growth, we treated mouse glioma cell lines (CT2A, GL261, and KR158) with the iron chelator deferoxamine (DFO) and the iron-donating compound ferric ammonium citrate (FAC), which inhibited and enhanced growth of these cells, respectively (**Figure 1B**). HFE is a critical iron regulator whose expression has been reported to correlate with GBM patient survival, although no mechanistic description has yet emerged to explain these observations^21^. We therefore investigated the role of HFE in GBM by first determining whether *HFE* levels in GBM patient tumors differ from those in non-tumor brain tissue. To do this, we utilized the GEPIA database^22^ and found significantly elevated *HFE* expression in GBM compared to non-tumor brain tissue (**Figure 1C**). Expression data from The Cancer Genome Atlas (TCGA) as well as the Chinese Glioma Genome Atlas (CGGA) revealed a direct correlation between *HFE* and tumor grade, with the highest expression seen in GBM (grade IV) (**Figure 1C**). Using these datasets, we also assessed the relationship between *HFE* levels and patient survival, with “high” and “low” *HFE* defined as the top and bottom 50% of expression. Although no differences were observed in male GBM patients, high *HFE* expression correlated with truncated survival in female patients in both datasets, confirming previous findings^21^ (**Figure 1D**). Importantly, high and low *HFE* expression was comparable in males and females, indicating that the differences observed are not based on sex-specific variations in *HFE* levels (**Supplemental Table 1**). As HFE primarily regulates iron status through modulation of other proteins, we also analyzed survival based on high and low expression of these iron-associated genes (**Supplemental Table 2**). Interestingly, high expression of ferritin subunits, ferritin heavy chain (*FTH*) and ferritin light chain (*FTL*), predicted poorer survival outcomes in female GBM patients, while no differences were present in male patients. Ferritin is responsible for intracellular iron storage and similar associations between ferritin expression and survival in GBM have previously been reported, as well as an inhibitory effect on tumor cell growth when ferritin expression is reduced^15^.These data suggest an important tumor-intrinsic role for HFE in GBM that may contribute to sex-specific effects on survival.

**Figure 1.**
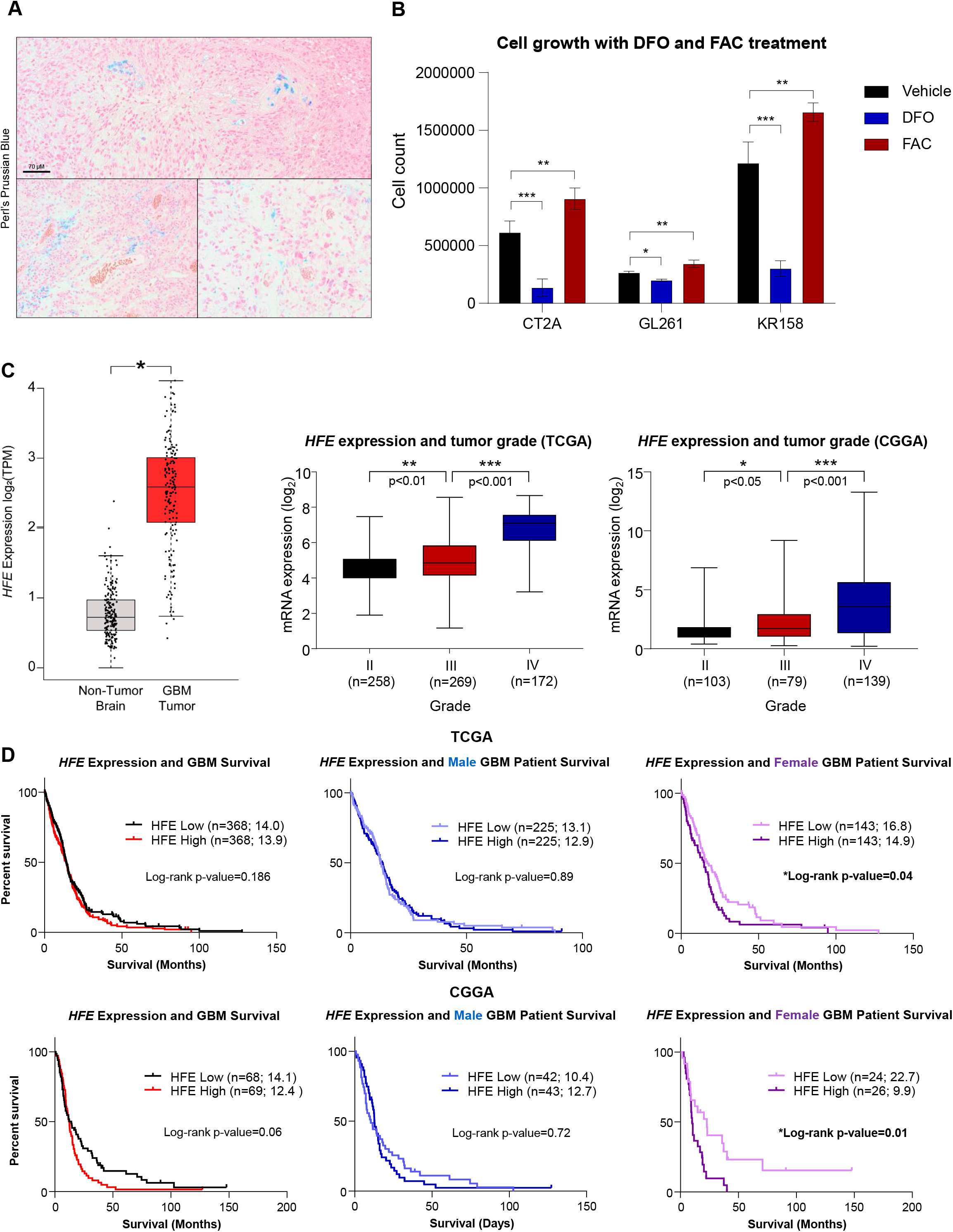
*HFE* is expressed in GBM and correlates with poorer survival in female patients. A) Perl’s Prussian blue stain to visualize iron content in human GBM tissue (scale bar at 70 μM). B) Cell growth measured by trypan blue exclusion in mouse glioma cells (CT2A, GL261, and KR158) after treatment with vehicle (water), the chelator deferoxamine (DFO, 10 μM) or iron donating agent ferric ammonium citrate (FAC, 15 μM). 50,000 cells were plated in triplicate in 6 well plates and collected and counted at days 1, 3, 5, and 7. C) *HFE* expression from GEPIA in GBM tumor (n=163) and non-tumor (n=207) brain tissue and *HFE* expression compared among glioma grades II-IV. D) Survival data from TCGA and CGGA comparing overall survival of male and female IDH-wildtype GBM patients based on high and low *HFE* expression (top and bottom 50% of expression). *p<0.05; **p<0.01; ***p<0.001 determined by t-test (first panel C), one-way ANOVA with Dunnett’s multiple comparisons test (B and second panel C), or log-rank test for survival data (D). Error bars represent standard deviation.

### Hfe knockdown in mouse glioma cells induces apoptosis in vitro and inhibits tumor growth in vivo

While *HFE* expression has been reported to inform GBM survival, its function in tumor cells remains unclear^21^. To directly assess its function, we genetically manipulated *Hfe* levels in tumor cells to determine the effect on cell growth and survival. We first measured baseline *Hfe* mRNA levels in three mouse glioma cell lines (CT2A, GL261, and KR158) compared to wild-type mouse astrocytes (**Figure 2A**). CT2A *Hfe* expression was significantly lower than that of astrocytes, while KR158 had significantly higher baseline expression. Based on these results, we utilized two separate, non-overlapping shRNA constructs to perform genetic knockdown of *Hfe* in KR158 cells. Knockdown 1 (KD1) and knockdown 2 (KD2) reduced *Hfe* mRNA expression by approximately 60% and 80%, respectively, compared to a non-targeting shRNA, and a reduction in protein expression was validated by immunoblot (**Figure 2B**). To determine the impact of *Hfe* knockdown on tumor cell growth *in vitro*, we first performed a trypan blue exclusion assay. By day 3 of growth, significantly fewer cells were present in both knockdown groups compared to the control, and caspase 3/7 activity, a surrogate of apoptosis, was significantly increased in both knockdown cell lines, suggesting greater induction of cell death with *Hfe* reduction (**Figure 2C**). To directly quantify proliferation, a dye dilution assay was performed in which cells were incubated with carboxyfluorescein succinimidyl ester (CFSE), a fluorescent dye that stains cells, and grown in culture. The cells were then collected, and fluorescence was quantified. Rapidly proliferating cells have a lower concentration of dye as it becomes diluted with each subsequent division, while slowly dividing cells have a higher dye concentration (**Supplemental Figure 1**). Quantification of CFSE dilution was significantly elevated with *Hfe* knockdown, indicating that the decreased cell number may also be driven by a decrease in the rate of proliferation (**Supplemental Figure 1**). Based on the observation that elevated *HFE* expression in female GBM patients is predictive of poorer survival, we intracranially implanted control and *Hfe*-knockdown cells into male and female C57Bl/6J mice to determine whether *Hfe* loss extends survival. We found increased tumor latency with *Hfe* knockdown in both male and female animals (**Figure 2D**). Interestingly, three of the female mice implanted with KD2 cells failed to present with symptoms after 100 days, at which point exploratory necropsy failed to reveal the presence of a tumor. Taken together, these data suggest that loss of *Hfe* induces an apoptotic phenotype in mouse glioma cells and slows tumor growth *in vivo*.

**Figure 2.**
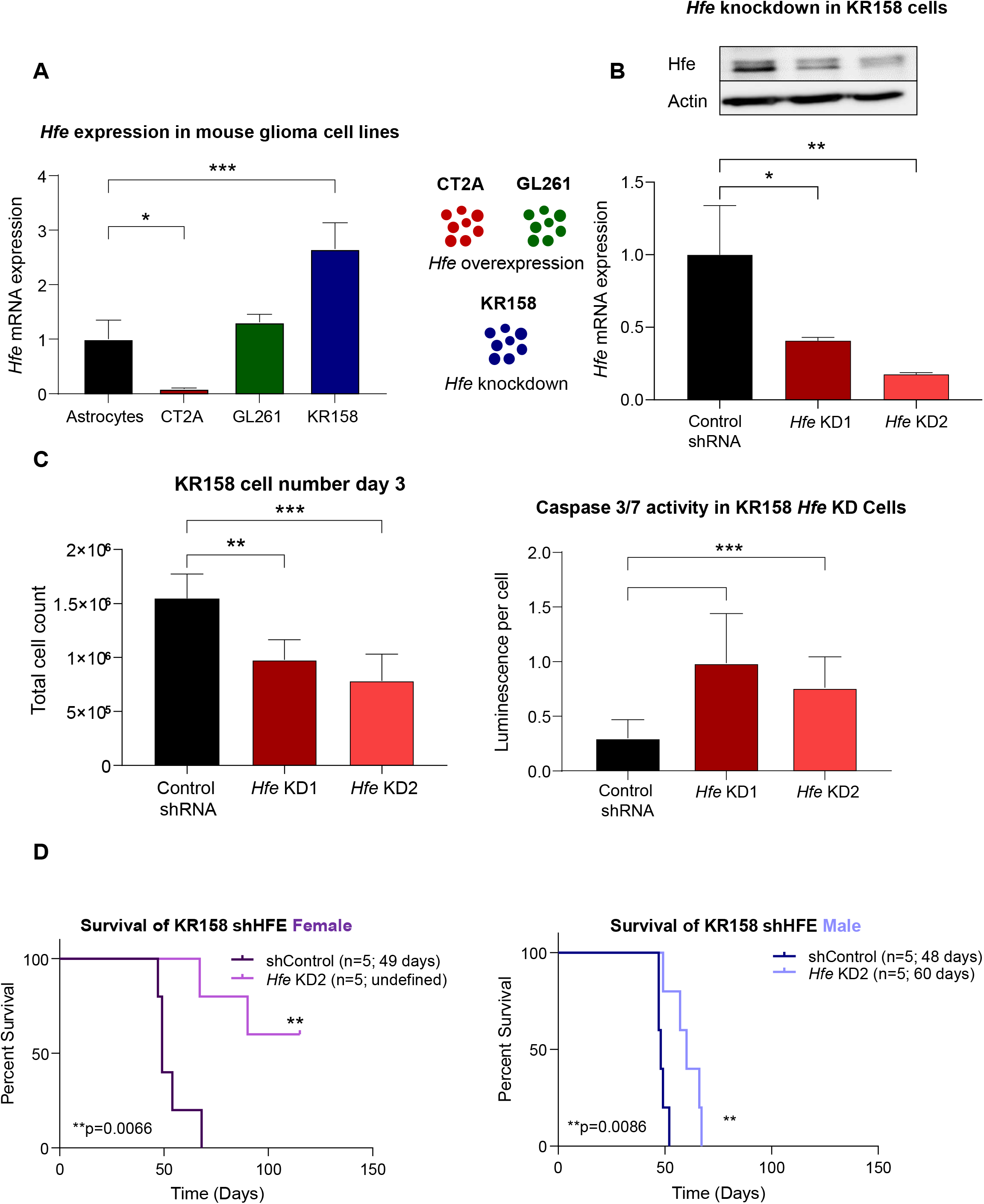
*Hfe* knockdown in KR158 cells induces apoptosis and significantly extends survival in female mice. A) RT-qPCR quantification of *Hfe* in mouse glioma cell lines (CT2A, GL261, and KR158) and wild-type mouse astrocytes, normalized to *Gapdh*. Based on expression levels, *Hfe* was knocked down in KR158 cells and overexpressed in CT2A and GL261 cells. B) RT-qPCR and western blot validation of *Hfe* knockdown in KR158 cells with actin as a loading control. C) Trypan blue exclusion was used to determine cell number in KR158 control and knockdown cells after 3 days of growth. Caspase 3/7 activity was measured after 3 days using Caspase-Glo and normalized to cell number. D) KR158-knockdown and control cells were intracranially implanted in male and female mice, with n=5 for each group, and survival was measured. Median survival is provided on plots. *p<0.05; **p<0.01; ***p<0.001 determined by one-way ANOVA with Dunnett’s multiple comparisons test (A-C) or log-rank test for survival data (D). Error bars represent standard deviation.

### Increased Hfe expression in mouse glioma cells drives proliferation and tumor growth in vivo

Based on the observation that CT2A and GL261 cells express lower and comparable levels of *Hfe*, respectively, compared to wild-type astrocytes, we performed genetic overexpression of *Hfe* in these cells using stable lentiviral transfection. Overexpression was validated by RT-qPCR and immunoblot in both cell lines (**Figure 3A**). Increased cell number as assessed by trypan blue exclusion was observed at day 5, with no difference in caspase activity (**Figure 3B, Supplemental Figure 2A**). To determine whether enhanced proliferation driven by *Hfe* overexpression could account for the differences in cell number, we assayed cell division by measuring CFSE dye diltuion. *Hfe* overexpression in both CT2A and GL261 cells significantly increased the proliferation rate compared to controls (**Supplemental Figure 2B**). Female mice implanted with both overexpression cell lines succumbed to disease more quickly than those implanted with control cells, while males exhibited no difference in survival outcomes with overexpression, suggesting that tumor growth is augmented only in females when *Hfe* levels are high (**Figure 3C**). These findings are consistent with the human GBM data and reveal a role for HFE in tumor cell proliferation and survival in females.

**Figure 3.**
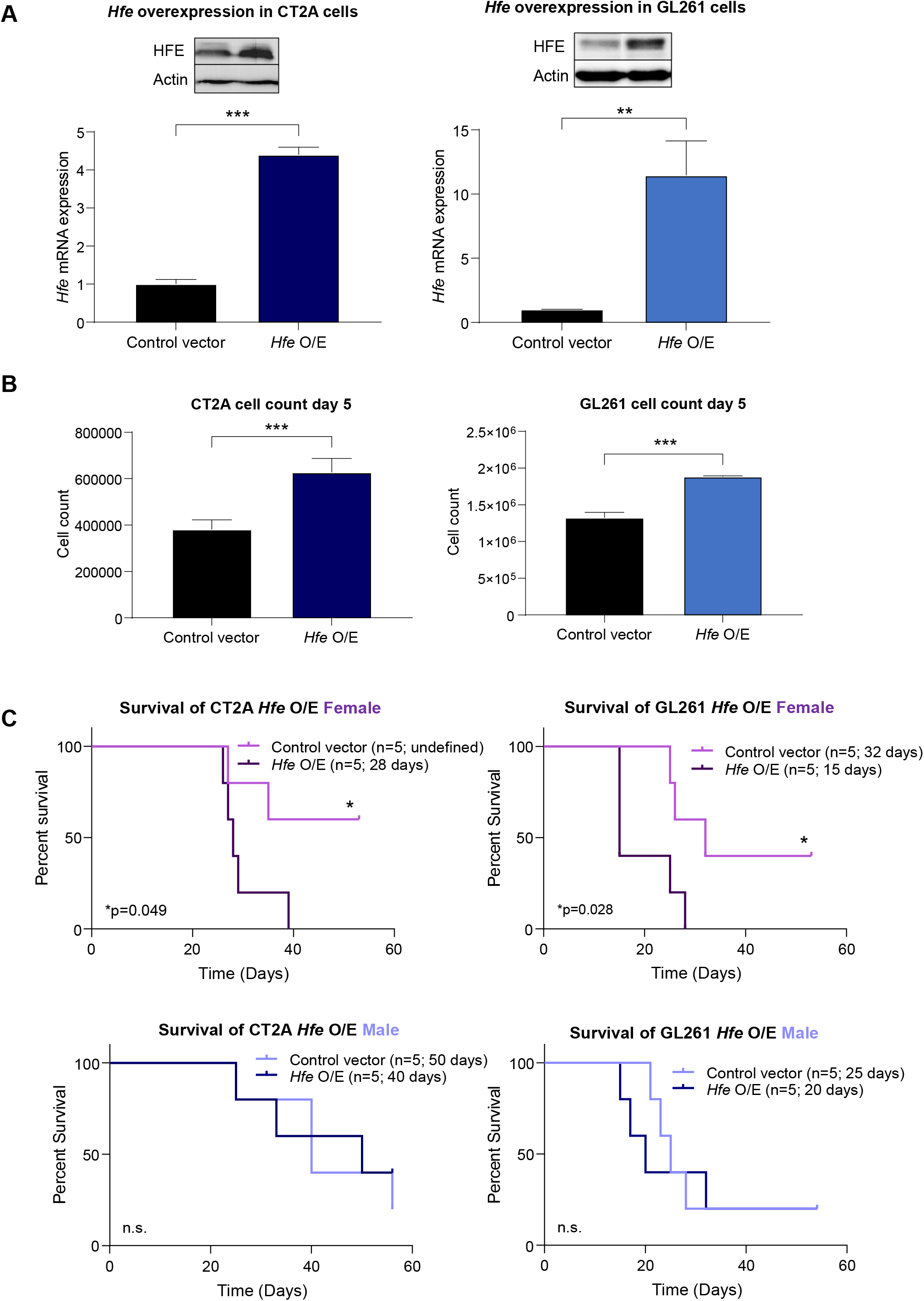
*Hfe* overexpression in CT2A and GL261 cells increases proliferation and significantly reduces survival in female mice. A) RT-qPCR and western blot validation of *Hfe* overexpression in CT2A and GL261 cells with actin as a loading control. B) Trypan blue exclusion was used to determine cell number in CT2A and GL261 control and *Hfe* overexpression cells after 5 days of growth. C) Intracranial implantation of CT2A and GL261 *Hfe* overexpression and control cells in male and female mice. *p<0.05; **p<0.01; ***p<0.001 determined by t-test (A and B) or log-rank test for survival data (C). Error bars represent standard deviation.

### Hfe knockdown increases iron uptake and the production of reactive oxygen species

Based on the role of HFE in metabolic iron regulation, we investigated GBM iron uptake using radioactive iron assays, which revealed significantly increased uptake in *Hfe* knockdown cells compared to control (**Figure 4A**). This is consistent with the known function of HFE as a competitive inhibitor of iron uptake through binding of transferrin receptor 1 (TFRC) on the cell surface. Overexpression of *Hfe* had no effect on iron uptake (**Supplemental Figure 3A**). As HFE does not directly interact with iron but rather with other iron-handling proteins, we hypothesized that modulating *Hfe* expression in these cells would in turn disrupt the expression of other iron-associated genes. We measured mRNA levels and observed a significant reduction in the expression of ferritin heavy chain (*Fth1*) and transferrin receptor 1 (*Tfrc*) and significant upregulation of ferroportin (*Slc40a1*) in knockdown cells compared to control (**Figure 4B**). Downregulation of transferrin receptor, which is involved in iron import, and upregulation of ferroportin, which is responsible for iron export, may represent an attempt by *Hfe*-knockdown cells to reduce high intracellular iron levels caused by increased uptake. Furthermore, a reduction in ferritin suggests a limited ability of these cells to store extra iron, which could result in cytotoxicity and induce apoptotic cell death. Overexpression of *Hfe* did not impact expression of transferrin receptor or ferritin, although ferroportin expression was significantly decreased in *Hfe* overexpression cells (**Supplemental Figure 3B**). To investigate mechanisms through which *Hfe* loss induces an apoptotic phenotype, we interrogated signaling pathway alterations after *Hfe* knockdown using the NanoString sequencing platform. A number of pathways were differentially expressed when comparing control cells to *Hfe* KD1 and KD2 (**Figure 4C**). Among these, oxidative stress was reported as the top pathway upregulated in both knockdown cell lines compared to control. Given our findings that *Hfe* loss induces a cell death phenotype, we measured the generation of reactive oxygen species (ROS) in these cells and saw a significant induction of hydrogen peroxide (H_2_O_2_) production with knockdown compared to control (**Figure 4D**). H_2_O_2_-mediated apoptosis is well documented and has can act through activation of caspases^23,24^. Collectively, these findings suggest that loss of *Hfe* function in KR158 cells results in an iron overload phenotype and increased production of ROS, leading to cell death.

**Figure 4.**
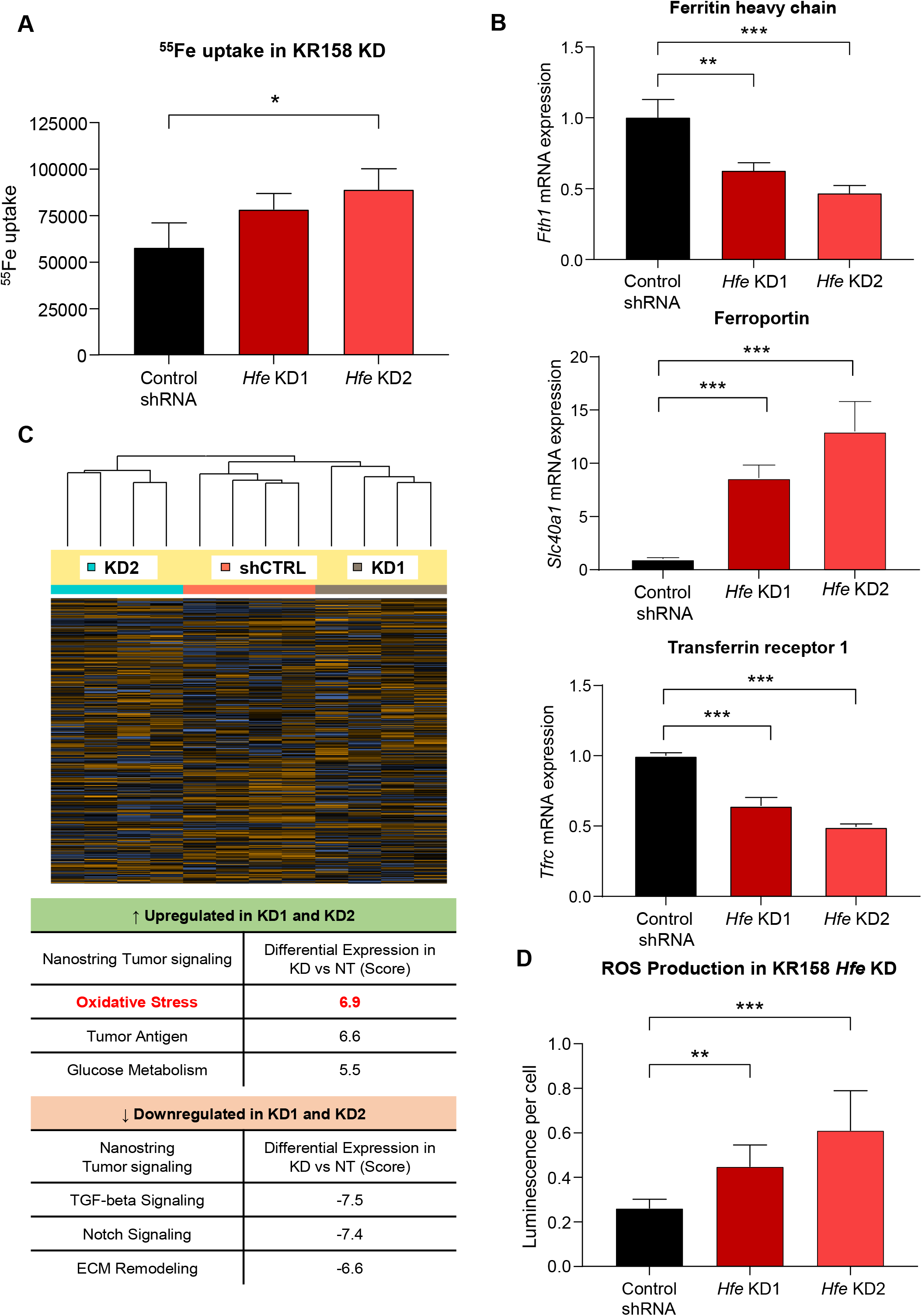
Loss of *Hfe* enhances iron uptake and induces the formation of reactive oxygen species (ROS). A) Radioactive ^55^Fe uptake normalized to total protein content. B) Iron-associated gene expression measured by RT-qPCR. Fold change compared to control and normalized to *Gapdh* is shown. C) NanoString overall clustering and pathway score comparing *Hfe* knockdown to control in KR158 cells. D) H_2_O_2_ production in *Hfe* knockdown and control cells was measured by ROS-Glo and normalized to cell number. *p<0.05; **p<0.01; ***p<0.001 determined by one-way ANOVA with Dunnett’s multiple comparisons test. Error bars represent standard deviation.

## Discussion

Tumor cells co-opt iron metabolic programs to satisfy their high iron demand and support rapid proliferation. Targeting iron metabolism has emerged as a popular strategy for the development of clinical trials in a number of different cancers, including brain, prostate, colon, liver, and lung^11^. These trials typically attempt to use chelating agents to reduce tumor iron availability, but their lack of specificity for tumor cells and promiscuity for other off-target divalent metal ions can lead to dose-limiting toxicities and diminish efficacy^25–28^. Thus, a better understanding of the molecular mechanisms governing aberrant tumor iron metabolism is necessary to improve therapeutic development. GBM tumors exhibit increased iron uptake, and metabolic iron regulation in this disease has primarily been investigated from the perspective of proteins directly involved in iron handling^15,29–31^. As an iron sensor, HFE is critical for the maintenance of intracellular iron homeostasis but indirectly regulates iron flux by binding and modulating expression of other iron-associated proteins. Previous work has described an association between increased *HFE* expression and truncated GBM patient survival^21^. Accordingly, we find that *HFE* is upregulated in GBM tumors compared to non-tumor brain tissue. However, the mechanisms underlying these observations have yet to be elucidated.

Iron uptake and storage can be exploited by tumor cells to enhance iron sequestration and promote tumor growth. GBM cancer stem-like cells demonstrate increased expression of iron uptake proteins transferrin (TF) and transferrin receptor (TFRC), as well as the iron storage protein ferritin^15^. Knocking down either subunit of ferritin (FTH1 or FTL) in these cells was sufficient to reduce tumorsphere formation *in vitro* and tumor progression *in vivo*^15^. Our findings demonstrate increased iron uptake with *Hfe* knockdown and a counterintuitive induction in apoptotic cell death. When we interrogated the impact of *Hfe* knockdown on iron-associated gene expression, we found a reduction in ferritin heavy chain (*Fth1*). Ferritin is essential for limiting the accumulation of intracellular ferrous iron (Fe^2+^), which can be used in the Fenton reaction to catalyze the formation superoxide and hydroxyl radicals (Fe^2+^ + H2O2 → Fe3+ + HO^•^ + OH^−^)^32^. These free radicals can cause oxidative stress and induce several forms of cell death, including caspase-dependent apoptosis^33^. Indeed, we found that *Hfe* knockdown induced ROS production in the form of H_2_O_2_, which is consistent with previous reports that ferritin degradation in human GBM cells promotes ROS generation and induces ferroptosis, an iron-dependent form of cell death^16^. In fact, ferroptotic induction via delivery of iron oxide nanoparticles has been reported as an effective therapeutic strategy in animal models of GBM^34^. This work demonstrated that increasing intracellular iron levels while simultaneously augmenting H_2_O_2_ production results in potent ROS generation and ferroptosis with minimal off-target toxicity^34^. Thus, a shift in focus from iron chelation, which broadly targets iron and other metal ions, to iron storage may represent a less toxic strategy in the treatment of GBM tumors (**Figure 5**).

**Figure 5.**
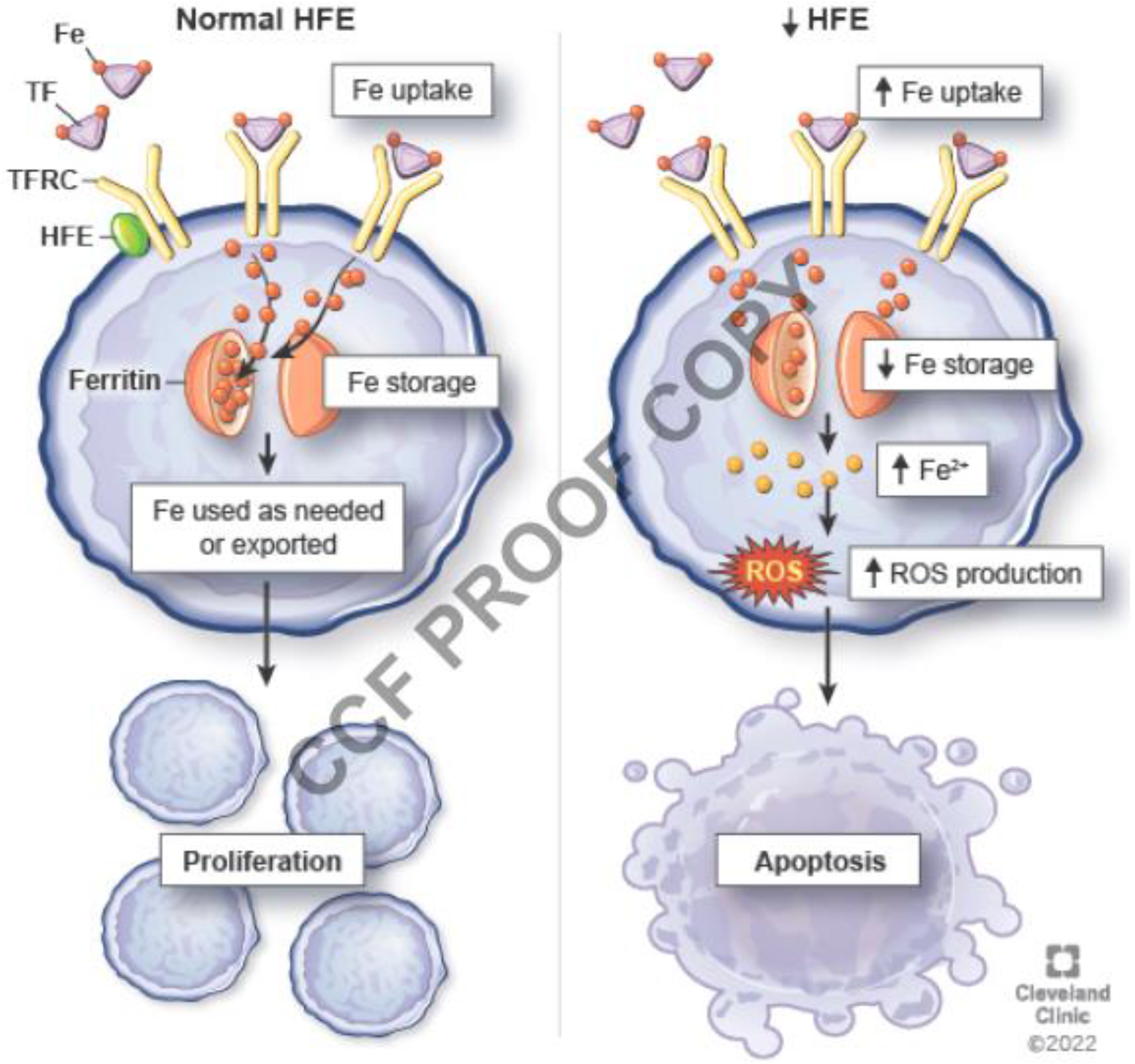
Loss of tumor cell intrinsic *Hfe* induces cell death through ROS generation. Under normal conditions (left panel) HFE associates with transferrin receptor (TFRC) on the cell membrane to competitively inhibit transferrin (TF)-bound iron uptake and maintain homeostasis. Our findings indicate that loss of HFE (right panel) permits greater iron uptake and results in downregulation of ferritin, leading to reduced iron storage, greater production of ROS, and an apoptotic phenotype.

Sex is an important determinant of tumor risk and survival outcomes in GBM^3^. Men are more likely to develop GBM at a ratio of 1.6:1 compared to females, and male GBM patients are reported to have worse prognoses, with an estimated median survival of 15.9 months compared to 22.6 months in female patients^3^. Our work supports previous findings that tumor-intrinsic HFE exerts a sex-specific effect on overall survival, with high HFE expression ablating the survival advantage normally observed in female patients^21^. The basis for these differences is still unclear but may be attributed to sex-mediated differences in iron metabolism. Men typically have larger iron stores than women, as much as two- to three-fold higher in some tissues, and women experience iron deficiency and anemia at much higher rates than men^35,36^. Historically, this has largely been attributed to blood loss through menstruation and conditions such as pregnancy, but other factors such as hormones likely play a role. For example, there is an inverse relationship between serum ferritin and estrogen levels which cause iron stores to gradually increase after menopause^37^. However, both serum ferritin and iron stores are still reportedly lower in post-menopausal women than in men^38,39^. A consequence of these sex differences in iron metabolism is that high serum iron levels are much more prevalent in men than women and have been associated with increased cancer risk^40^. We observed truncated survival only in female mice implanted with *Hfe*-overexpressing tumor cells. This finding is consistent with human GBM clinical data and may be partially explained by sex differences in iron levels. Conversely, *Hfe* knockdown led to protracted survival in both male and female animals. We suspect that this survival benefit is mediated by the apoptotic phenotype and may therefore not discriminate based on sex. Taken together, these results suggest a role for sexual dimorphism in HFE-mediated tumor iron regulation that ultimately results in the observed differential survival benefits.

While these findings are one of the first descriptions of HFE function in GBM, this work has limitations. Due to the complex regulatory nature of HFE, it is difficult to determine the direct target and sequence of events leading to apoptosis upon *Hfe* depletion. It is possible that these effects are mediated primarily by changes in iron status with HFE modulation. However, HFE also forms cell surface protein complexes that initiate downstream signaling and may directly influence ROS production or apoptosis. Further complicating our understanding of this process is the regulation of iron-associated gene and protein expression by iron availability. The mRNA transcripts of most iron-associated genes possess iron response elements on their 5’ UTRs, which regulate translation rates. Thus, HFE may modulate expression directly, through intracellular signaling, or indirectly through iron availability, and more insight into this complex relationship is necessary to fully appreciate its role in tumor cells. Another limitation is the asexual state of our mouse tumor cell lines. Upon karyotyping, we discovered that these lines have only one X chromosome and the original sexes of the animals from which they were derived are unknown. Thus, our ability to thoroughly interrogate the role of tumor sex differences was limited. The development of new syngeneic mouse GBM cell lines, in which the cell sex is known, would be beneficial to fully appreciate these differences. Despite these limitations, HFE remains an intriguing mechanism for targeting tumor iron flux. In conclusion, our findings demonstrate that HFE drives tumor cell proliferation and may be a viable approach for targeting iron metabolism and inducing cell death.

## Materials and Methods

### Perl’s Prussian Blue staining

Formalin fixed paraffin embedded patient samples were assembled as tissue microarrays and kindly provided by Dr. Bjarne Kristensen with approval by the Regional Committee on Health Research Ethics for Southern Denmark (S-20150148). Staining of tumor sections with Perl’s Prussian blue and nuclear fast red counterstain was performed by the Lerner Research Institute Imaging Core.

### Patient mRNA expression and survival

Gene expression data from The Cancer Genome Atlas (TCGA; date accessed: 4/15/2021) and the Chinese Glioma Genome Atlas (CGGA; date accessed: 2/1/2021) were downloaded from the GlioVis data visualization portal^41^. Additional data including patient sex and IDH status were also obtained from Gliovis and IDH mutant samples were excluded from the analysis. High and low HFE were defined as the top and bottom 50% of expression.

### Cell culture

Syngeneic mouse GBM cell lines (CT2A, GL261, and KR158) were grown in adherent conditions in RPMI 1640 media with 10% fetal bovine serum (FBS) and 1% penicillin-streptomycin. Media was replaced every other day and cells were passaged with Accutase and phosphate buffered saline when sub-confluent (70-80%). Cells were maintained in humidified incubators at 37 °C and 5% CO_2_.

### Cell treatment with DFO and FAC

Deferoxamine (DFO; Sigma; D9533) was reconstituted in DMSO at a concentration of 76mM and further diluted to a 5mM stock solution in water. Ferric ammonium citrate (FAC; Sigma; RES20400-A7) was diluted in water at a concentration of 5mM. Cells were treated with 10 μM DFO or 15 μM FAC for 3 days prior to collection and counting.

### Cell viability

For trypan blue exclusion, cells were washed with PBS, trypsinized, collected, and spun down at 300xg for 5 minutes. Cells were then resuspended in media and 20 μL of cells were taken for counting. An equal volume of trypan blue (20 μL) was added to the cells and mixed thoroughly. 10 μL of the mixture was applied to a cell counting slide (Bio-Rad; 1450003) and measured using a Bio-Rad Automated Cell Counter.

### Immunoblot

Protein was isolated from cells using NP-40 lysis buffer consisting of 10 mM Tris HCl, 1 mM EDTA,150 mM NaCl, 0.5% NP-40, 1 mM PMSF, 1x protease inhibitor (Sigma; p8340), and 1x phosphatase inhibitor cocktail (Sigma; p5726). Cells were resuspended in buffer, lysed, and vortexed prior to being spun down for 30 minutes at 20,000 rpm. The supernatant was collected and protein concentrations were measured using protein assay dye (Bio-Rad; 5000001) with a bovine serum albumin (BSA) standard. Protein was diluted in SDS sample buffer and 30 μg of each sample was loaded per well into polyacrylamide SDS-PAGE gels. Gels were run at 120 volts for 110 minutes and transferred onto PVDF membranes (Millipore; IPVH00010). Membranes were blocked with 5% nonfat milk and probed with anti-HFE antibody (Santa Cruz; sc-514405) and β-Actin (Bio-Rad 12004163, 1:10,000) as a loading control. Horseradish peroxidase conjugated secondary antibodies were added to the membranes (anti-rabbit, Invitrogen; 31460 and anti-mouse EMD Millipore; AP308P). Membranes were developed with ECL Western Blotting Substrate (Thermo Scientific; 32106) and read with a Bio-Rad ChemiDoc Imaging System.

### Quantitative real-time PCR

RNA was extracted from cells using an RNeasy kit (Qiagen; 74134) and concentrations were measured using a NanoDrop spectrophotometer. cDNA was synthesized using qScript cDNA SuperMix (Quanta Biosciences; 101414-102). qPCR was performed in Fast SYBR Green Mastermix (Applied Biosystems; 01120793) and an Applied Biosystems QuantStudio 3. Primer sequences are shown in Supplemental Table 3. During qPCR analysis, threshold cycle values were normalized to *Gapdh*.

### HFE knockdown and overexpression

MISSION® pLKO.1-puro Non-Mammalian shRNA Control Plasmid (SHC002) and HFE shRNA plasmids TRCN0000105417 (KD1), TRCN0000105419 (KD2) were purchased from Sigma. 293T cells were used to package lentivirus with psPAX2 and pMD2G using calcium phosphate transfection. Lentiviral particles were collected from media and concentrated using a PEGit virus precipitation solution. For transfection, cells were grown in 10cm tissue culture plates and concentrated lentivirus was added to cells. After 24 hours of incubation, media was changed and incubated for an additional 24 hours prior to selection with puromycin (5 μg/mL).

### Intracranial tumor implantation

Intracranial implantation experiments with syngeneic tumor cell lines were performed as previously described^42^. 6-week-old C57Bl/6J mice were anesthetized using inhaled isoflurane and an insulin syringe attached to a stereotaxic apparatus was used to inject cells into the left hemisphere at a depth of approximately 3.5mm. Each syringe was prepared with equal cell numbers suspended in 10μL of null RPMI 1640 media (20,000 KR158 cells transfected with shcontrol or *Hfe* KD2; 10,000 CT2A or GL261 cells transfected with control vector or *Hfe* overexpression). Animals were monitored over time for the presentation neurological and behavioral symptoms associated with end-point. Investigators were blinded to experimental conditions while monitoring animals. All animal experiments were performed in compliance with institutional guidelines and were approved by the Institutional Animal Care and Use Committee of the Cleveland Clinic (protocol 2019-2195).

### Statistical analysis

For two-group comparisons, p-values were calculated using unpaired t-test. For multiple group comparisons, one-way ANOVA with Dunnett’s multiple comparisons test was used. Log-rank tests were used for *in vivo* survival analysis. All statistical analyses were performed using GraphPad Prism 6. All *in vitro* experiments were carried out in at least technical triplicate for each experimental group. Additional statistical details, including p values and sample size can be found in figure legends.

## Acknowledgements

The authors thank all members of the Lathia and Connor laboratories for thoughtful discussion and project conceptualization. We thank Drs. Trine Jorgensen, Erin Murphy, Jessica Williams, and Jennifer Yu for their invaluable feedback with regard to this work. We also thank P01 collaborators Drs. Jill Barnholtz-Sloan, Michael Berens, and Joshua Rubin for their intellectual contributions to this work. We are grateful to Ms. Karen Keslar and Dr. Robert Fairchild for their technical assistance with Nanostring analysis and to Ms. Amanda Mendelsohn from the Center for Medical Art and Photography for illustrations.

## Notes

**Disclosure of Conflicts of Interest:** No conflicts of interest to declare.

### Competing Interest Statement

The authors have declared no competing interest.

