## Supplementary Material for "Tumor cell-intrinsic HFE drives glioblastoma growth"

**Supplemental Figure Legends**

**Supplemental Figure 1.**
A) CFSE is used to measure proliferation. During each round of cell division, intracellular CFSE becomes diluted and is quantified to determine proliferation rate. B) Flow cytometry plots of CFSE dilution in KR158 *Hfe* knockdown cells compared to control.

**Supplemental Figure 2.**
A) Flow cytometry plots of CFSE dilution in CT2A and GL261 *Hfe* overexpression cells. B) Caspase 3/7 activity measured using Caspase-Glo and normalized to cell number.

**Supplemental Figure 3.**
A) Radioactive $^{55}Fe$ uptake normalized to total protein content. B) Iron-associated gene expression measured by RT-qPCR. Fold change compared to control is shown.

**Supplemental Table 1. TCGA and CGGA *HFE* expression levels in male and female GBM patients.**

**Supplemental Table 2. TCGA and CGGA overall survival in male and female GBM patients with high and low expression of iron-associated genes.**

**Supplemental Table 3. Mouse RT-qPCR primers used in this study.**

Supplemental Figure 1

**A**

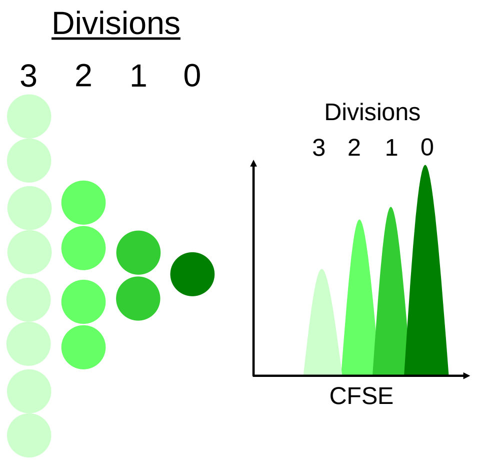


**B**

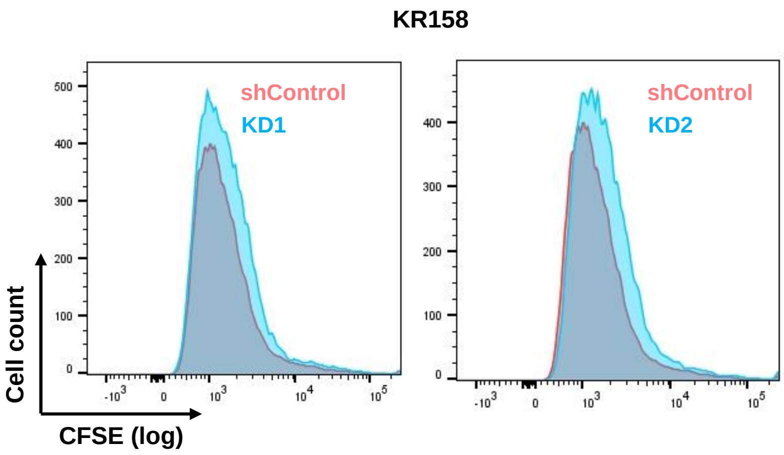


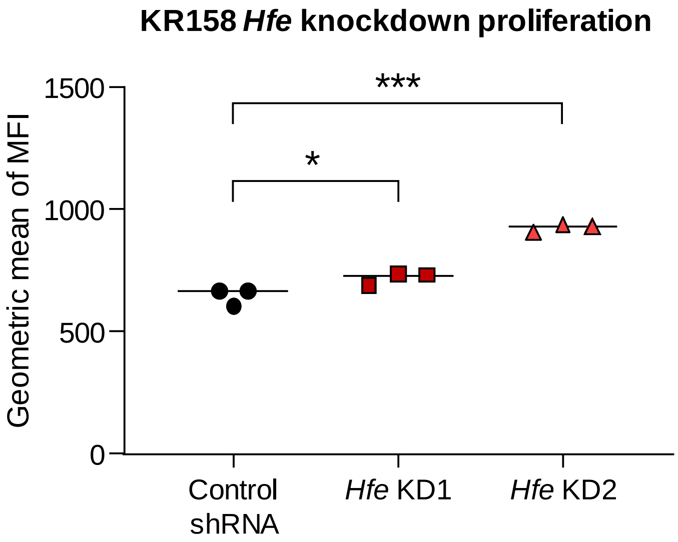

Supplemental Figure 2

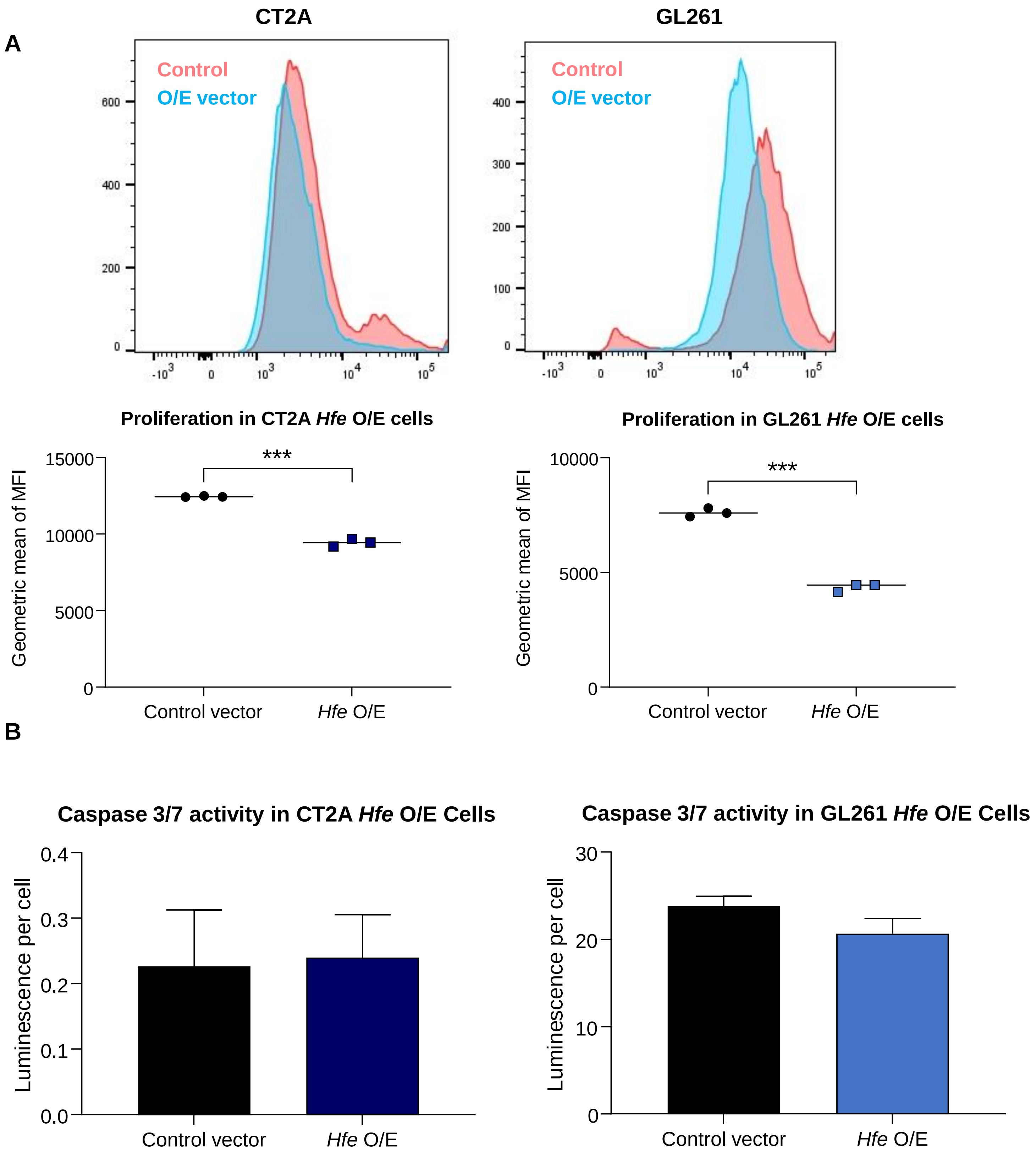

Supplemental Figure 3

A

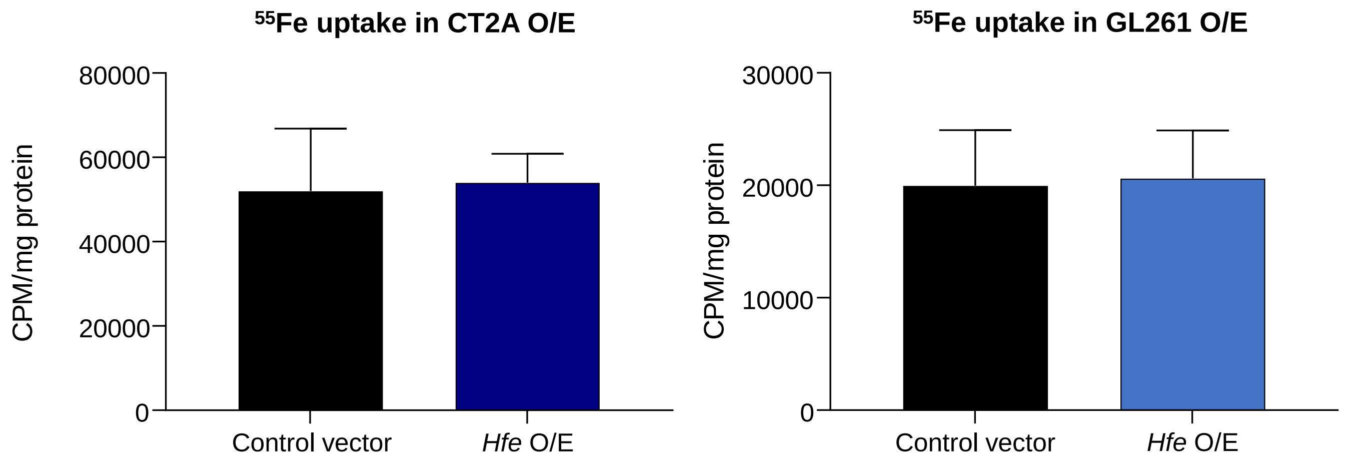


B

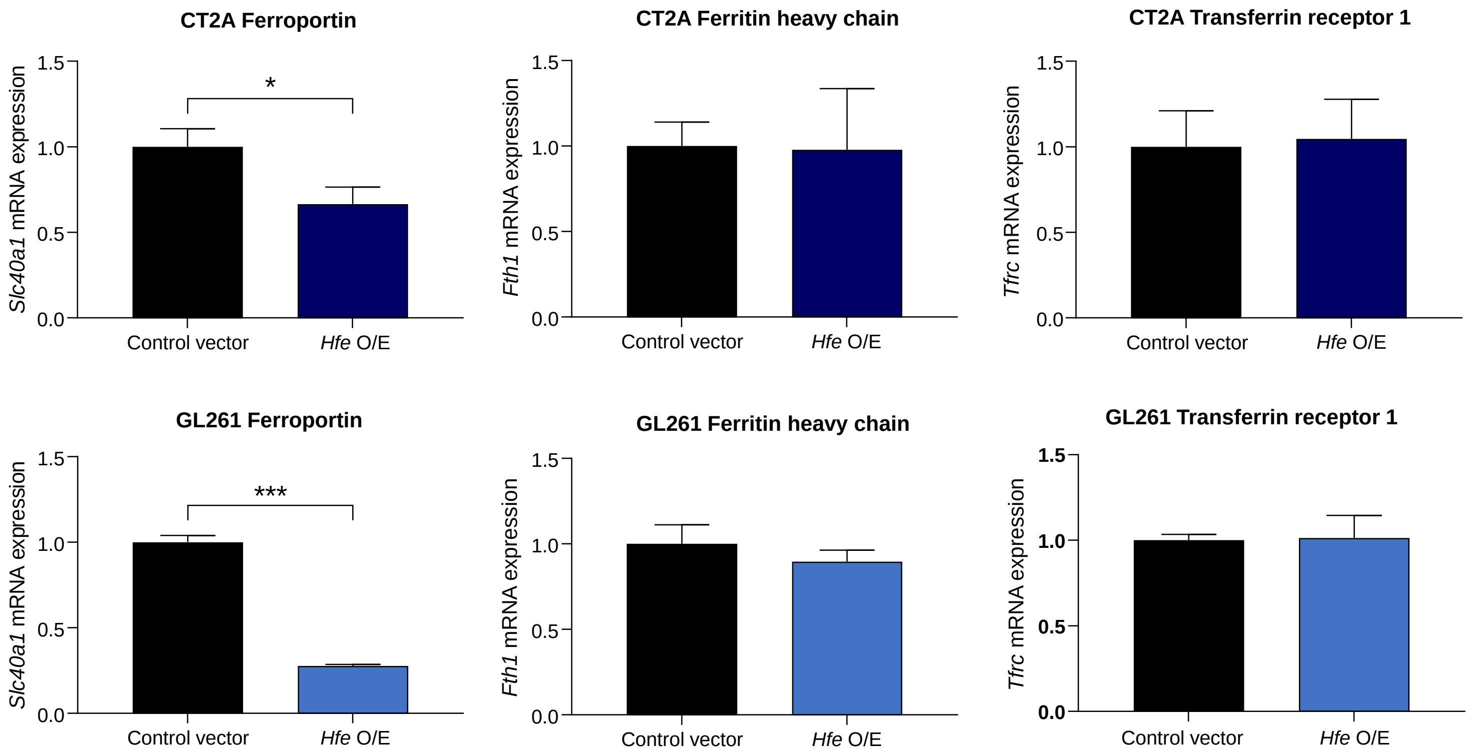

Supplemental Table 1

| *HFE* expression TCGA ($log_2$) | | |
|---|---|---|
| Sex | Low | High |
| Male | 4.06 | 4.38 |
| Female | 4.06 | 4.45 |
| *HFE* expression CGGA ($log_2$) | | |
| Sex | Low | High |
| Male | 0.83 | 2.53 |
| Female | 0.59 | 2.19 |

Supplemental Table 2

| All patients (n=368) | | | |
|---|---|---|---|
| Gene | Median survival months (Low) | Median survival months (High) | P-value |
| *FTH* | 14.9 | 12.7 | **<0.05*** |
| *FTL* | 15.0 | 12.2 | **<0.05*** |
| *HAMP* | 14.1 | 13.9 | 0.82 |
| *HFE* | 14.0 | 13.9 | 0.19 |
| *TFRC* | 13.9 | 13.9 | 0.82 |
| Female patients (n=143) | | | |
| Gene | Median survival months (Low) | Median survival months (High) | P-value |
| *FTH* | 18.3 | 12.6 | **<0.05*** |
| *FTL* | 17.6 | 12.2 | **<0.05*** |
| *HAMP* | 15.0 | 16.0 | 0.66 |
| *HFE* | 16.8 | 14.9 | **<0.05*** |
| *TFRC* | 14.9 | 15.7 | 0.09 |
| Male patients (n=225) | | | |
| Gene | Median survival months (Low) | Median survival months (High) | P-value |
| *FTH* | 14.0 | 12.7 | 0.50 |
| *FTL* | 14.1 | 12.2 | 0.40 |
| *HAMP* | 13.3 | 12.9 | 0.35 |
| *HFE* | 13.1 | 12.9 | 0.89 |
| *TFRC* | 12.9 | 13.3 | 0.21 |

Supplemental Table 3

| **Gene** | **Forward** | **Reverse** |
|---|---|---|
| *Fth1* | CTCATGAGGAGAGGGAGCAT | GTGCACACTCCATTGCATTC |
| *Gapdh* | AACAGCAACTCCCACTCT TC | CCTGTTGCTGTAGCCGTATT |
| *Hfe* | CACCGCGTTCACATTCTCTA | AAAGAGCTGGTCATCCACATAG |
| *Slc40a1* | CGGTCTTTGGTCCTTTGATTTG | GCAGAAGGTCAAGAAGGTAGTT |
| *Tfrc* | AGCCAGATCAGCATTCTCTAAC | TCTGCAGCCAGTTTCATCTC |

### Supplemental Materials and Methods

*CFSE proliferation assay*

*Hfe* knockdown or overexpressing cells and respective controls were stained with 1 μmol/L carboxyfluorescein succinimidyl ester (CFSE; BioLegend 423801) for 15 minutes at 37°C and washed with twice with PBS. Stained cells were then plated at equal densities (50,000 KR158 and CT2A; 100,000 GL261) in 6 well plates and allowed to grow for 2 to 3 days. Samples were analyzed using a BD LSR Fortessa.

*Flow cytometry*

Samples were stained with LIVE/DEAD (Thermo Fisher Scientific; L34961) diluted at 1:500 in PBS for 10 minutes at room temperature. Cells were then washed once in PBS and resuspended in Mojosort Buffer (Biolegend; 480017) at a 1:5 dilution in dd$H_2O$. Samples were analyzed with a BD LSR Fortessa (BD Biosciences), and FlowJo (Version 10.5.0, FlowJo LLC) was used for data analysis. Compensation controls were used to account for spectral overlap.

*Radioactive iron uptake*

$^{55}Fe$ uptake was performed as previously described[43]. Cells were grown to 70-80% confluence, washed, and incubated in serum-free RPMI 1640 medium for 24 h. The cells were incubated with $^{55}Fe$-NTA in the same medium for 4 h at 37°C in a 5% $CO_2$ incubator. The medium was aspirated and the cells were washed twice with 150 μM NaCl 100 μm EDTA to remove excess iron. $^{55}Fe$-NTA uptake was measured in triplicate wells by lysis in RIPA buffer followed by liquid scintillation counting. All values were normalized to total protein concentration as determined by a Bradford assay.

*Nanostring*

RNA was isolated using an RNeasy mini kit (Qiagen) and the nCounter® Tumor360 Panel was subsequently used to analyze gene expression. Two non-overlapping shRNA *Hfe* constructs (KD1 and KD2) in KR15 cells were analyzed in triplicate. nSolver version 4.0 was used to normalize and analyze data to determine up- and downregulated pathways in both knockdown conditions.

*ROS quantification*

ROS production was quantified using the ROS-Glo $H_2O_2$ assay (Promega; G8820) according to the manufacturer's protocol. Cells were plated in triplicate in a 96 well plate at equal density (1,000 cells per well) and allowed to grow for 2 to 3 days prior to incubation with $H_2O_2$ substrate solution for 4 hours. The ROS-Glo detection solution was then added and incubated for 20 min. Luminescence was measured using a Victor3 plate reader (PerkinElmer) and values were normalized to cell number.
